# metabolisHMM: Phylogenomic analysis for exploration of microbial phylogenies and metabolic pathways

**DOI:** 10.1101/2019.12.20.884627

**Authors:** E.A. McDaniel, K. Anantharaman, K.D. McMahon

## Abstract

**Summary:** Advances in high-throughput sequencing technologies and bioinformatic pipelines have exponentially increased the amount of data that can be obtained from uncultivated microbial lineages inhabiting diverse ecosystems. Various annotation tools and databases currently exist for predicting the functional potential of sequenced genomes or microbial communities based upon sequence identity. However, intuitive, reproducible, and user-friendly tools for further exploring and visualizing functional guilds of microbial community metagenomic sequencing datasets remains lacking. Here, we present metabolisHMM, a series of workflows for visualizing the distribution of curated and user-provided Hidden Markov Models (HMMs) to understand metabolic characteristics and evolutionary histories of microbial lineages. metabolisHMM performs functional annotations with a set of curated or user-defined HMMs to 1) construct ribosomal protein and single marker gene phylogenies, 2) summarize the presence/absence of metabolic pathway markers, and 3) create heatmap visualizations of presence/absence summaries.

**Availability and Implementation:** metabolisHMM is freely available on Github at https://github.com/elizabethmcd/metabolisHMM and on PyPi at https://pypi.org/project/metabolisHMM/ under the GNU General Public License v3.0.

## 1. Introduction

Common comparative genomic approaches for analyzing large metagenomic datasets include analyzing the distribution and evolutionary history of genes of interest, describing the presence/absence of specific metabolic pathways in metagenome assembled genomes (MAGs) or single cell genomes (SAGs), and comparing these results to existing publicly available genomes. Many of these steps require computational expertise in several bioinformatic tools, specific file formats, and sometimes use of expensive, proprietary software platforms. Tools for intuitively summarizing and visualizing the functional potential of sequenced genomes in a high-throughput, user-friendly, reproducible manner that allow for maximum user flexibility (i.e. custom marker sets) and making comparisons among large genome datasets are overall lacking. Here we present metabolisHMM, a set of reproducible workflows for executing common comparative genomics analyses using profile Hidden Markov Models (HMMs). metabolisHMM encompasses a set of easy-to-use and flexible workflows for visualizing phylogenies and metabolic heatmaps from both curated and custom-provided HMM-based profile annotations. We demonstrate the capabilities and output results of metabolisHMM by using publicly available bacterial and archaeal genomes from a subsurface aquifer system (1), available as an online tutorial at https://github.com/elizabethmcd/metabolisHMM/wiki/Subsurface-Aquifer-Tutorial.

## 2. Implementation

The metabolisHMM package contains two main functionalities: 1) phylogeny construction to visualize evolutionary histories and 2) heatmap-based synthesis of metabolic pathway distributions. The basic input requirements for each of the four embedded workflows are DNA sequences as raw genomic scaffolds in fasta file format for subsequent gene prediction using Prodigal (2). Each workflow uses either curated or user-provided custom profile Hidden Markov Models (HMMs) with a threshold cutoff score provided by the user for performing functional annotation, based on the hmmsearch option of HMMER (3). Additionally, the user defines an output directory in which all intermediate files such as reformatted fasta files, HMM output results, alignments, and final phylogenies and heatmap figures are deposited. The remaining arguments and steps are workflow dependent. Detailed installation instructions and documentation for using the metabolisHMM package and each workflow is provided in the repository wiki: https://github.com/elizabethmcd/metabolisHMM/wiki.

### 2.1 Constructing single marker phylogenies

The single-marker-phylogeny workflow searches a panel of genomes for a specific gene marker and builds a phylogenetic tree. Any of the package-provided curated marker sets or a user-provided marker can be used for constructing a single-marker phylogeny. The alignment is constructed using MAFFT (4), and the user can choose to construct the phylogenetic tree using either FastTree (5) or RAxML (6), depending on available computational resources. Due to common issues with MAG gene content redundancy and unknown consequences of copy number variation from uncultivated organisms, metabolisHMM only uses the top-scoring hit for a particular marker within a genome for constructing the final alignment and phylogeny. Given a corresponding metadata file, the user can output data files configured for viewing trees with the interactive Tree Of Life (iTOL) online tool (7).

### 2.2 Creating genome phylogenies

The create-genome-phylogeny workflow takes a set of input genomes and creates a ribosomal phylogeny or species tree. We provide a set of 16 single copy ribosomal proteins as part of the metabolisHMM software package release that are specific for archaea or bacteria (32 markers total) as described in Hug et al. (8). Alignments and tree construction are performed as described above, with individual alignments concatenated across all genomes. Since metabolisHMM was developed specifically for comparing MAGs and SAGs against isolate genomes, metabolisHMM will warn the user if a genome contains less than 12 or a pre-defined value of ribosomal markers, as confidence in the phylogenetic reconstruction will be low if a genome is missing several markers in the final alignment, due to incompleteness.

### 2.3 Summarizing broad metabolic features using curated and custom markers

The summarize-metabolism workflow uses a set of manually curated profiles spanning major transformations in the carbon, nitrogen, sulfur, and hydrogen cycles, that were constructed and made publicly available by Anantharaman et al. (1). Marker descriptions are provided in the ancillary data files of the software distribution. The user also provides a metadata file containing either the specific taxonomical names for each genome, or broad groups by which to aggregate sets of genomes together, such as by phylum-level placement or sample origin. Any marker-genome pair with a value greater than 1 is changed to a value of 1, resulting in a table of 0’s and 1’s for the absence and presence, respectively, of every marker-genome pair. The resulting heatmap shows the presence/absence of all markers spanning broad biogeochemical cycles to show the overall functional guilds of the input genomes. In addition to visualizing curated marker sets provided with the metabolisHMM package, the user can specify any marker sets that are custom-made or from outside databases, such as the PFAM and TIGRFAM databases, and/or the recently released KofamKOALA distribution. (9). The search-custom-markers workflow takes a set of specified markers in a user-provided order and produces a heatmap similar to that of the broad summaries mentioned above.

## 3. Results and Assessment

To demonstrate the main features of the metabolisHMM workflow, we used a set of 2545 publicly available bacterial and archaeal genomes from an aquifer metagenomic dataset (1). All demo figures are available within the tutorial at https://github.com/elizabethmcd/metabolisHMM/wiki/Subsurface-Aquifer-Tutorial. Using the single-marker-phylogeny workflow, we created a phylogeny of the folD marker, part of the reductive acetyl-coA pathway (10). We created a corresponding ribosomal phylogeny of genomes containing the folD marker. Using the FastTree option for constructing phylogenies, to search all 2,545 genomes for the folD marker, construct the phylogeny of the single marker, and make a corresponding ribosomal phylogeny of the 610 hits was completed in less than 30 minutes using 1 thread on a standard laptop (2015 MacBook Pro). We then characterized the broad metabolic capabilities of a subset of groups of MAGs within the aquifer dataset. Genomes were aggregated by phylum or superphylum group, where the shade of the cell for a specific marker indicates the percentage of genomes within that group that contain each marker. To screen the 874 genomes for all 80 curated markers, this workflow completed in approximately 1 hour using 1 thread. Using the search-custom-markers workflow, we screened 874 genomes for the main steps and subunits that are part of the methyl and carbonyl branches of the reductive acetyl-CoA cycle (11). Markers were accessed from the KofamKOALA KEGG distribution of HMMs and the corresponding threshold cutoffs for each marker was used as suggested (9). For screening 874 genomes with 15 markers this workflow completed in less than 30 minutes using 1 thread.

We compared the main functionalities and unique capabilities included in metabolisHMM with other recently released and popular software pipelines used for visualizing various functional aspects of sequenced genomes (Table 1). This includes GtoTree, MetaSanity, METABOLIC, KEGG Decoder, and Anvi’o (12–16). Overall, the core functionalities of metabolisHMM are distributed among several existing pipelines. However, metabolisHMM allows for maximum flexibility concerning external HMM profiles, making a corresponding ribosomal phylogeny of genomes with a particular single marker, and customized groupings and orderings of heatmap visualizations. Additionally, metabolisHMM allows for all of these core functions and powerful customized options through simple workflows that are easy to install and use reproducibly.

**Table 1:** Comparison of metabolisHMM functionalities with other pipelines. If a particular software pipeline was not intended for certain functions, we denoted that with NA. Pipelines encompassing a functionality but does not include flexible or customizable options, for example, are denoted with an X. Packages with a comparable functionality to metabolisHMM are denotated with a ✔.

| Tool | Phylogenies of single markers | Phylogenomics/ phylogenies | Corresponding ribosomal tree of single marker hits | Curated metabolic markers/summaries | Custom marker input options | Heatmap visualizations | Custom group aggregating/row ordering |
| --- | --- | --- | --- | --- | --- | --- | --- |
| GtoTree | X | ✓ | X | NA | ✓ | NA | NA |
| MetaSanity | NA | ✓ | NA | ✓ | ✓ | ✓ | ✓ |
| METABOLIC | NA | NA | NA | ✓ | X | X | X |
| KEGG Decoder | NA | NA | NA | ✓ | X | ✓ | X |
| Anvi'o | ✓ | ✓ | X | NA | ✓ | ✓ | ✓ |
| metabolisHMM | ✓ | ✓ | ✓ | ✓ | ✓ | ✓ | ✓ |

## Acknowledgements

We would like to thank members of the McMahon lab for testing and providing feedback on the metabolisHMM package. We would like to specifically thank Dr. Sarah Stevens of the Data Science Hub at the University of Wisconsin – Madison for early feedback and troubleshooting.

